## Supplementary Figures for "BEEM-Static: Accurate inference of ecological interactions from cross-sectional metagenomic data"

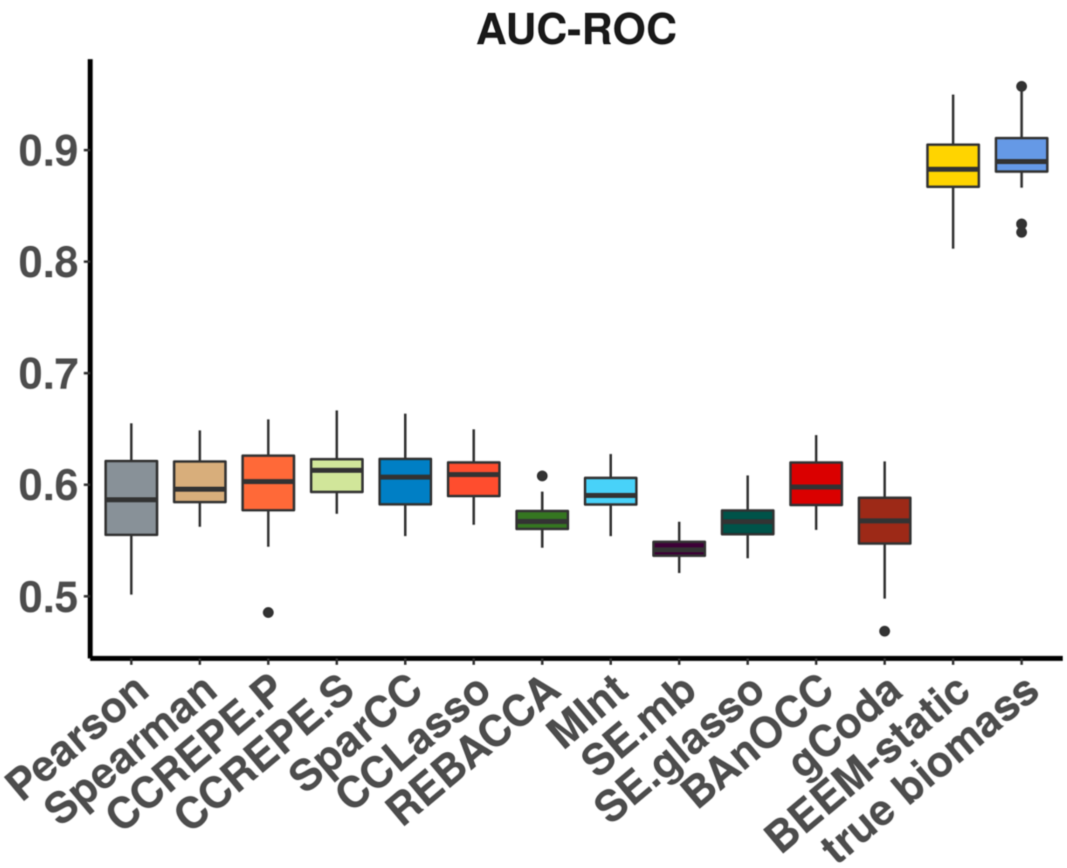


**Supplementary Figure 1. Performance comparison for determining interaction network structure.** Boxplots show AUC-ROC values for 30 different simulated communities with 30 species and 500 samples each.

A

B

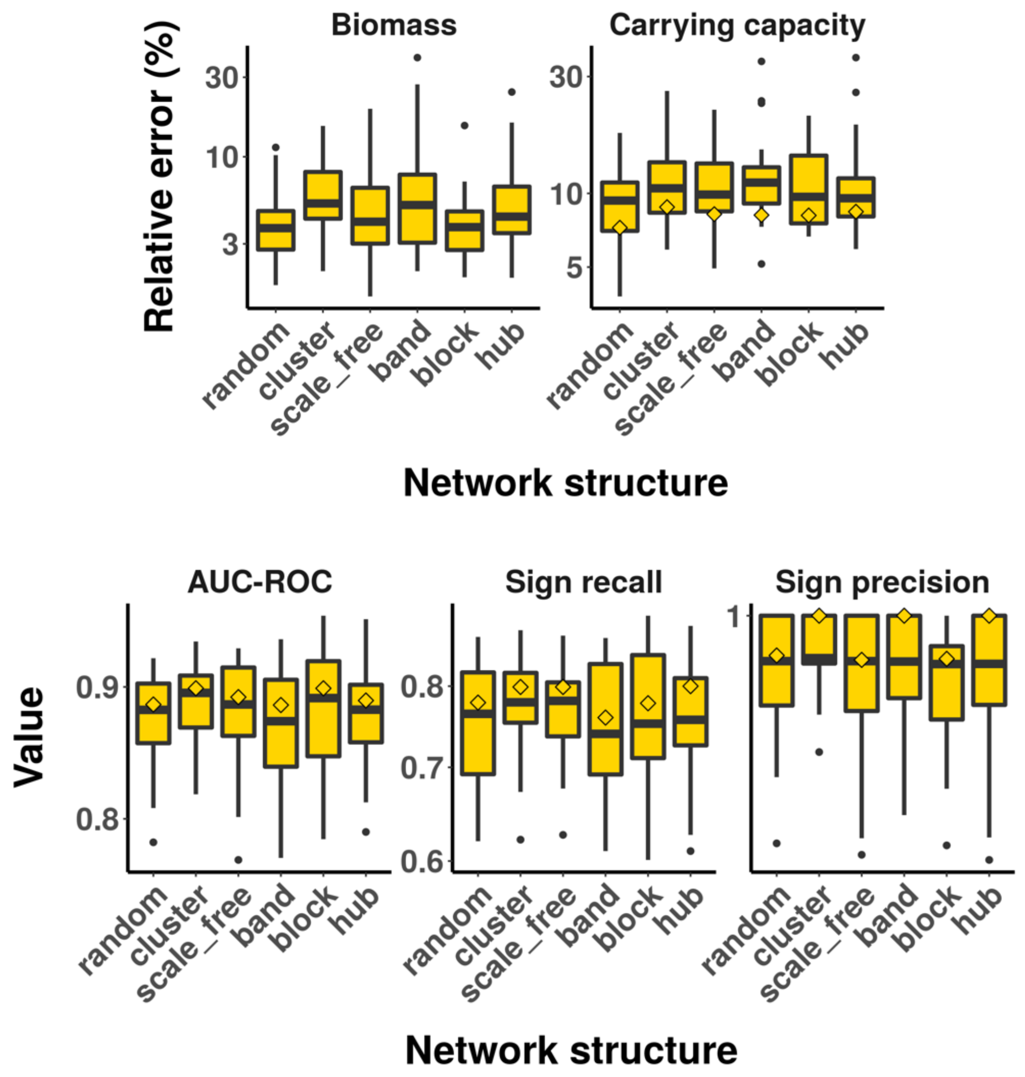


**Supplementary Figure 2. Robustness of BEEM-Static performance across different network structures.** Boxplots showing relative error for (A) parameter estimates for biomass (scaled to have the same median as the ground truth) and carrying capacity, and (B) interaction network AUC-ROC as well as sign recall and precision, based on 30 simulations. Diamonds mark median performance using true biomass values. Different network structures (except “random”) were generated with the SPIEC-EASI R package.

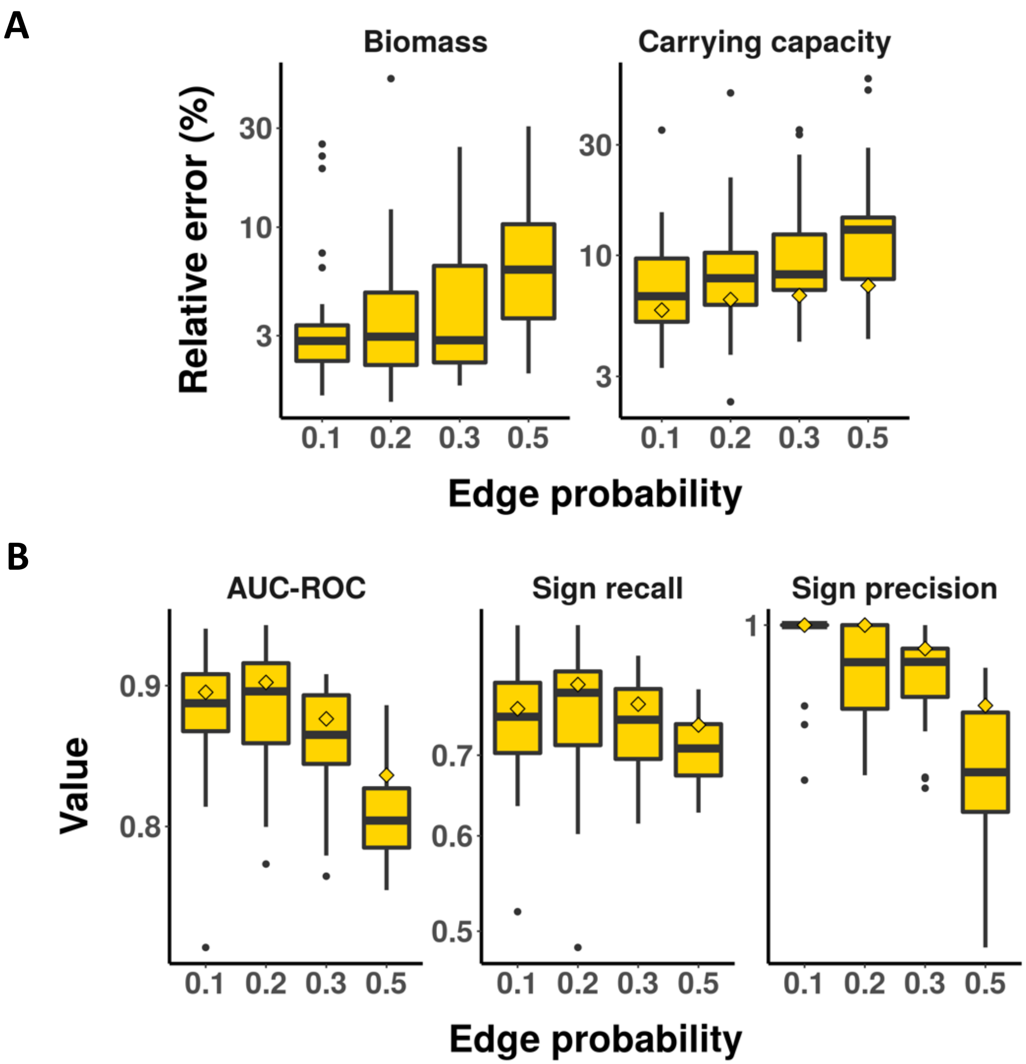


**Supplementary Figure 2. Robustness of BEEM-Static performance with varying edge densities.** Boxplots showing relative error for (A) parameter estimates for biomass (scaled to have the same median as the ground truth) and carrying capacity, and (B) interaction network AUC-ROC as well as sign recall and precision, based on 30 simulations. Diamonds mark median performance using true biomass values.

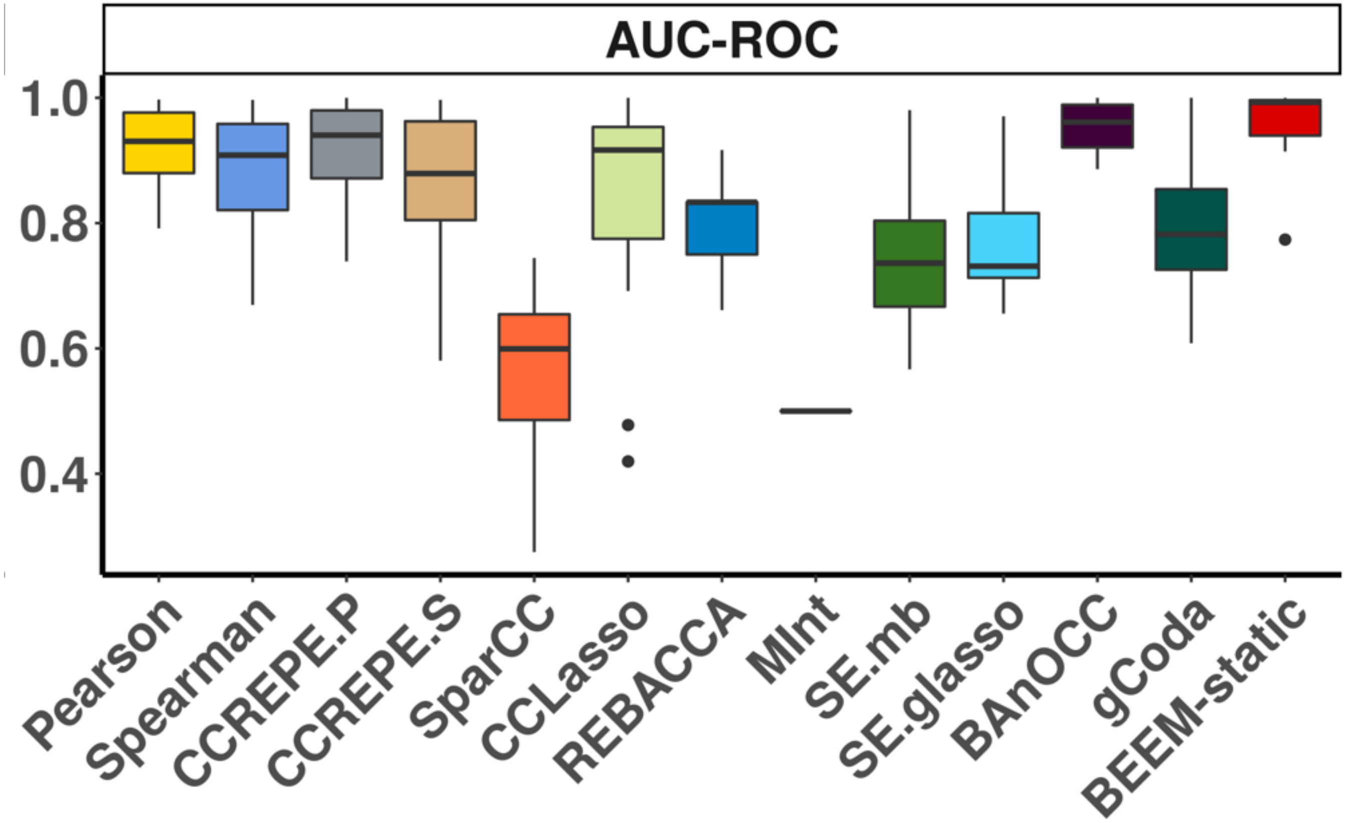


**Supplementary Figure 4. Benchmarking on synthetic communities created based on co-growth experimental data.** Boxplots show the performance of various methods based on 30 replicates that use randomly selected interacting pairs.

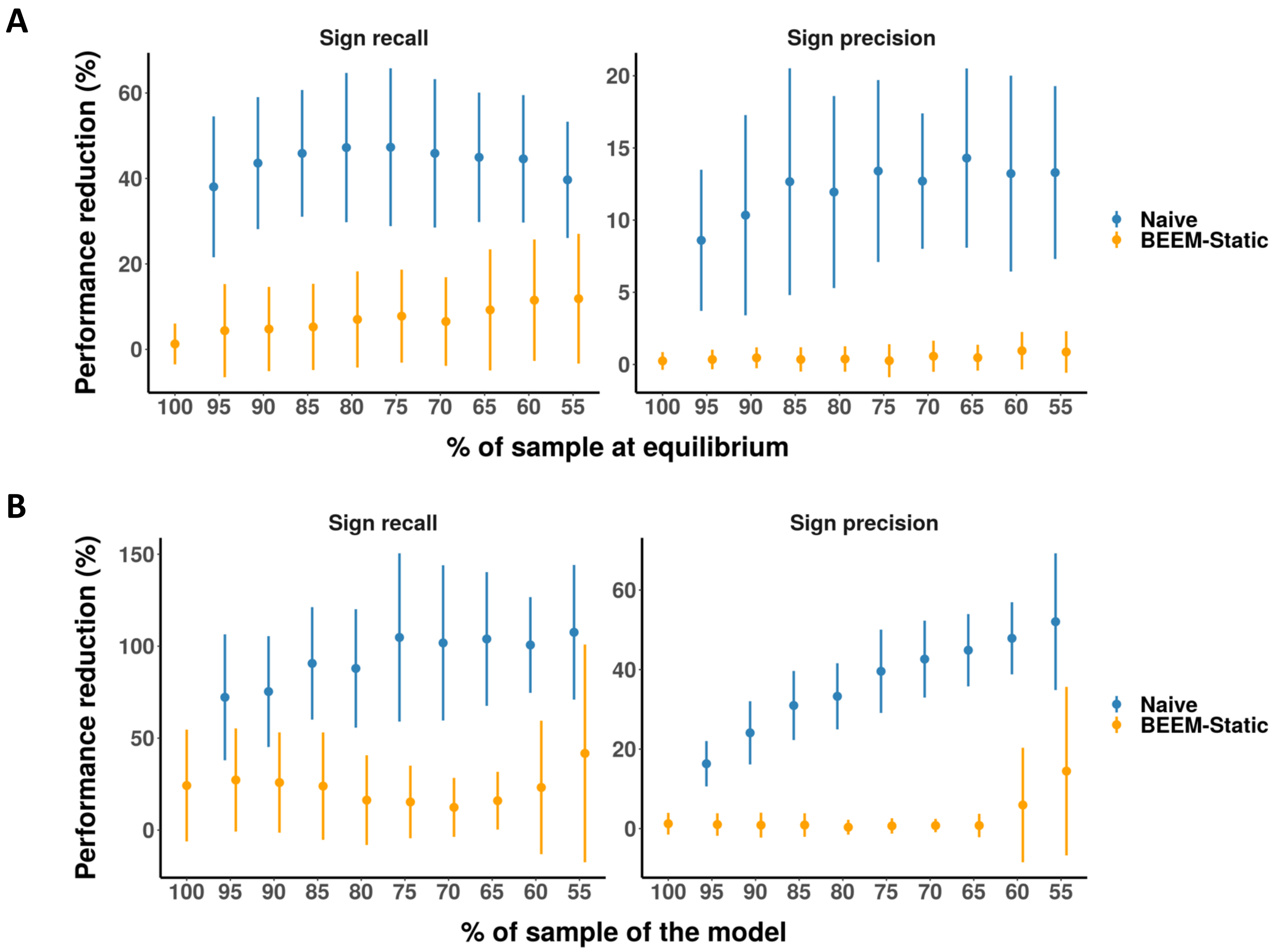


**Supplementary Figure 5. Utility of BEEM-Static filters for avoiding performance reduction in the presence of model violations.** Results are shown for increasing proportion of samples that are, (A) not at equilibrium or (B) not from the main model. The naïve algorithm is without filtering of samples and performance reduction is measured relative to BEEM-Static with no model violations for the data. Points show means while error bars show standard deviation across 30 simulations.

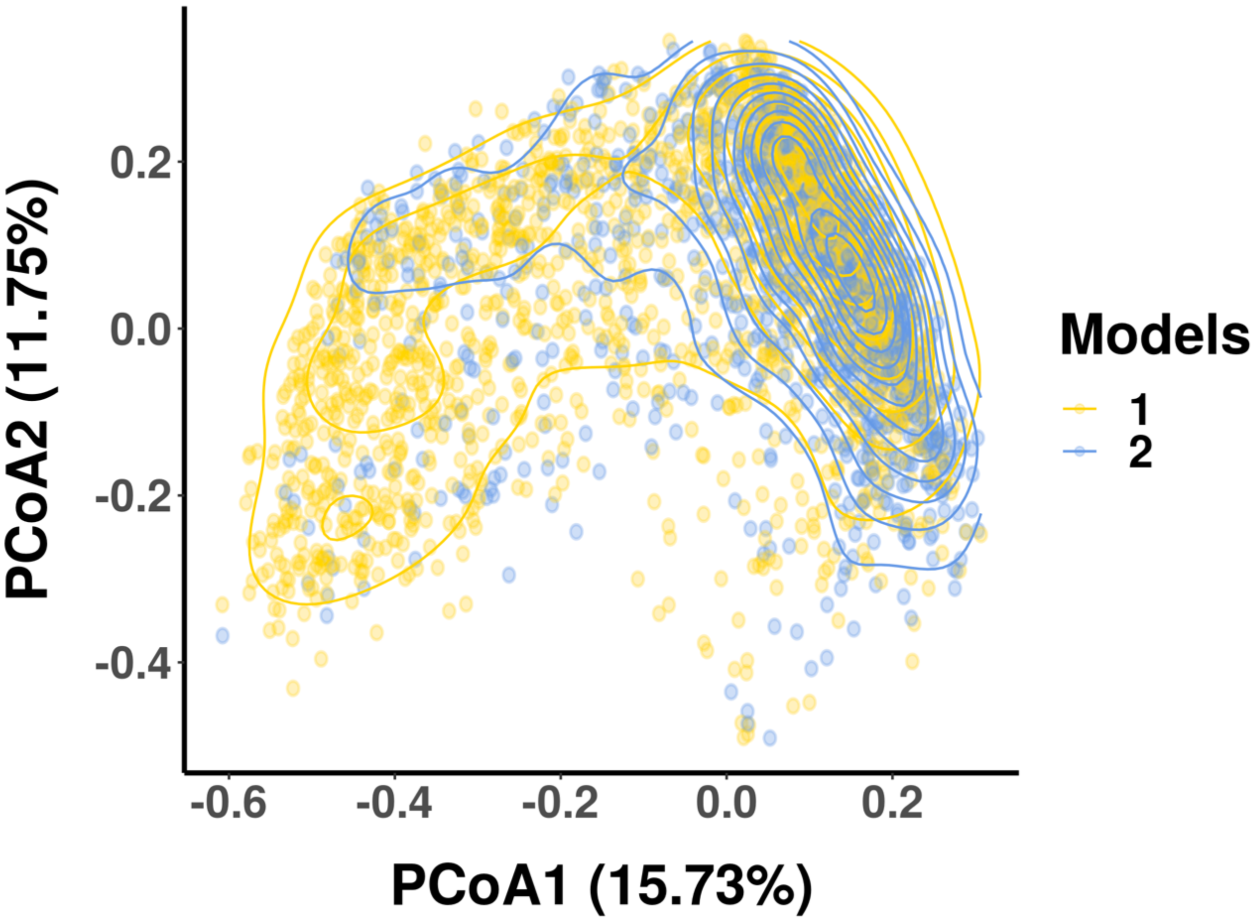


**Supplementary Figure 6. Ecological models from BEEM-static represent distinct configurations from enterotypes**. Principle coordinates plot (Bray-Curtis dissimilarity) based on gut microbiome taxonomic profiles. The contour lines highlight the two different regions where points aggregate. Points are colored by the BEEM-static models that they belong to (unassigned samples not shown), showing that while there appears to be some enrichment, model ids and enterotypes do not show a 1-1 correspondence.
